# *Serratia sp*. dominates the lung microbiome of patients with tuberculosis and non-tuberculous mycobacterial lung diseases

**DOI:** 10.1101/2024.05.27.596016

**Authors:** Meriem Belheouane, Barbara Kalsdorf, Stefan Niemann, Karoline I. Gaede, Christoph Lange, Jan Heyckendorf, Matthias Merker

**Author notes:** Correspondence to: Prof. Matthias Merker, Research Center Borstel, Leibniz Lung Center, Parkallee 1, 23845 Borstel. former affiliation (Jan Heyckendorf).

## Abstract

**Background:** Pathogenic mycobacteria, such as the *Mycobacterium tuberculosis* complex (Mtbc), and non-tuberculous mycobacteria (NTMs) can cause severe chronic pulmonary infections. However, not all infected patients develop active disease. Yet, it is unclear whether certain key taxa in the lung microbiome play a role in the pathogenesis of tuberculosis (TB) and NTM lung disease (LD).

**Material and methods:** We employed 16S rRNA amplicon sequencing (V3-V4) to characterize the baseline microbiome in bronchoalveolar lavage fluid from a patient cohort diagnosed with TB (n=23), NTM-LD (n=19), or non-infectious disease (n=4) prior to the initiation of therapy. The analysis included the depletion of human cells, removal of extracellular DNA, implementation of a decontamination strategy, and exploratory whole-metagenome sequencing (WMS) of selected specimens.

**Results:** The genera *Serratia* and unclassified *Yersiniaceae* dominated the lung microbiome of all patients with a mean relative abundance of >15% and >70%, respectively. However, at the sub-genus level, as determined by amplicon sequence variants (ASVs), TB-patients exhibited increased community diversity, and TB specific ASV_7 (unclassified *Yersiniaceae*), and ASV_21 (*Serratia*) signatures. Exploratory analysis by WMS and ASV similarity analysis suggested the presence of *Serratia liquefaciens*, *Serratia grimesii*, *Serratia myotis* and/or *Serratia quinivorans* in both TB and NTM-LD patients. Overall, presence/absence of certain *Serratia* ASVs was significantly associated with disease state.

**Conclusion:** The lung microbiome of TB patients harbors a distinct, and heterogenous microbiome structure with specific occurrences of certain *Serratia* traits. *Serratia sp.* plays a pivotal role in our understanding of microbial interactions in the lung microbiome of patients infected with Mtbc.

## Introduction

Mycobacteria belong to a genus with more than 200 species. Only few species can cause severe respiratory infections. *Mycobacterium tuberculosis* complex (Mtbc) bacteria cause tuberculosis (TB), a leading cause of morbidity and mortality worldwide [1]. Although the WHO estimates that approximately a quarter of the global population has been infected with Mtbc, only a minor fraction develops active TB. The main risk factors for developing TB are prolonged contacts to active pulmonary TB patients, immunodeficiencies (*e.g.* HIV-infection), diabetes mellitus, malnutrition, and tobacco use [1]. Non-tuberculous mycobacteria (NTMs), such as *M. avium-complex* or *M. abscessus* can cause NTM lung disease (LD). Transmission of NTMs occurs mostly via environmental sources, direct human-to-human transmission is very rare. Predominant risk factors for NTM-LD are previous pulmonary TB, cystic fibrosis, and bronchiectasis [2, 3]. However, the impact of commensal microbes colonizing the lower airways, *i.e*. the lung microbiome, on pathogenesis of TB and NTM-LD is often neglected.

Few studies have started to address the critical role of the gut and lung microbiome in the onset, progression, and susceptibility to mycobacterial lung diseases [4–7]. Indeed, healthy lungs are not sterile. Landmark studies established the biogeography of the healthy human lung microbiome, and revealed that the upper and lower airways harbor distinct microbiome compositions, while the lower compartment has a reduced microbial biomass [8–10]. Besides, Dickson and colleagues [11] validated bronchoscopy as a reliable method for investigating the lung microbiome and demonstrated that bronchoalveolar lavage fluid (BALF) specimens generate a consistent description of the resident taxa and are less prone to internal and external contamination. Although BALFs generate an adequate picture of the lung microbiome [12], these low-biomass specimens require thorough expertise in microbiome data generation and subsequent analyses [13]

To achieve a better understanding of the lung microbiome in TB and NTM-LD, we investigated BALF specimens collected in a retrospective cohort study at the Research Center Borstel, Germany, over 14 years. In comparison to BALF specimens from patients with non-infectious inflammatory lung diseases, we sought to identify key taxa associated with TB and NTM-LD.

## Materials and methods

### Study design

BALF was routinely collected from patients at the Medical Clinic of the Research Center Borstel (Germany) between July 2007 and September 2021 as part of diagnostic procedures in individuals with radiological abnormalities on thoracic imaging and symptoms of mycobacterial diseases, including fever, night-sweats, weight loss and cough, if mycobacterial DNA had not been detected genotypically in three sputa. The final diagnosis was reached on the basis of BALF microbiology and/or cytology, and radiology results. Bronchoscopy with BALF was performed on these patients according to national guidelines with 200 ml normal saline in fractions of 20 ml each [14]. Sterilizing procedures were performed following national guidelines [15]. For this study, we retrospectively included 63 bio-banked specimens from patients with a confirmed diagnosis of TB, NTM-LD, or non-infectious inflammatory lung disease. Previous medications and secondary diagnoses were retrospectively retrieved from the patient records.

### 16S rRNA sequencing and processing

Detailed protocols for depletion of host cells and extracellular DNA, as well as 16S rRNA sequencing of the V3-V4 hypervariable region are provided in the supplementary material. DNA libraries were prepared with a one-step PCR approach using 30 cycles, and sequenced on a MiSeq v3 kit with 2x300 bp paired-end reads. We included negative extraction controls, a dilution series of a microbial cell standard, pure cultures from different species, and technical replicates. The resulting fastq files were first processed with dada2 R package (v.1.16.0) [16], then subjected to a thorough multi-step decontamination scheme, and adjustment of clustering thresholds (supplementary material). Finally, sequencing depth was normalized to 10,000 reads for each sample, adjusted ASVs were further clustered into 98% OTUs, and 97% OTUs (supplementary material, and supplementary table S1).

### Ecological and statistical analyses

Statistical analyses were carried out in R (v. 4.2.1) [17]. The mean relative abundances of the main phyla, genera, adjusted ASVs, and 98% OTUs were compared across patient groups using the non-parametric Kruskal‒Wallis test. To identify indicator taxa of disease states, we applied indicator value analysis, calculated confidence intervals of the indicator value components, and evaluated the coverage of the identified indicator taxa among disease states in indicspecies” R package (v.1.7.13) [18] (supplementary material)

To explore the interactions between different taxa within patient groups, we calculated Spearmańs correlation between the relative abundances of ASVs and 98% OTUs. To evaluate the diversity and community structure of the lung microbiome within and across patients, we calculated several diversity indices with “vegan” (v.2.6-4) [19] at the adjusted ASV and 98% levels. The within-individual diversity was evaluated by Shannon (observed diversity) and Chao1 (expected richness) indices, and compared across patient groups using the non-parametric Wilcoxon test. To examine community structure, and assess the effect of patient and sampling characteristics along with disease state, we first calculated the Bray‒Curtis and Jaccard indices, then applied the non-parametric ANOVA “adonis” with 10^5^ permutations, and constrained principal coordinates analysis. The tested variables were disease group, age group, sex, smoking status, steroid medication, and sampling year.

### Whole-metagenome sequencing (WGS)

We performed an exploratory metagenomic analysis to determine the *Serratia* species in BALF DNA extracted from patients with a concentration >1 ng/µL (NTM patients, n=3) (detailed methodology in supplementary material).

## Results

### Study cohort

We analyzed 46 out of 63 available BALF specimens which passed the required data quality, and depth. These included adult patients (18 years and older) with pulmonary TB (n=23), NTM-LD (n=19), and non-infectious inflammatory lung disease (n=4) (one sample per patient). To the best of our knowledge, specimens were taken prior to the therapy against the main diagnosis, and as part of the routine diagnosis during an inpatient stay between July 2007 and September 2021. Patient age categories spanned between 10-19 years to 80-89 years, with 60-69 and 70-79 comprising the majority of the patients, *i.e.*, 13 and 14 patients, respectively. Thirty-five percent (n=16) of the patients were female. Individual TB-patients harbored viral infections, namely, hepatitis and HIV, while some NTM patients were diagnosed with chronic obstructive pulmonary disease (COPD), bronchitis, or pneumonia. Detailed patientś characteristics are provided in supplementary table S2.

### Composition of the lung microbiome in patients with pulmonary TB and NTM-LD

To assess the similarities, and disparities of the lung microbiome across patients with pulmonary TB, with NTM-LD and patients with other pulmonary diseases, we compared the most abundant taxa, and found that *Proteobacteria* strongly and similarly dominated the lungs of all three patient groups (figure 1a, supplementary figure S1a). In addition, we observed no significant differences in the abundances of the major phyla among the patient groups (Kruskal‒Wallis test, p > 0.05). At the genus level, the genera “unclassified *Yersiniaceae”* and *Serratia* (which also belongs to the family *Yersiniaceae*) dominated the lungs of all three patient groups (mean relative abundances of >70%, and >15%, respectively, while the genus *Mycobacterium* was detected in only one NTM patient (figure 1b). Few individuals were highly dominated by a single taxon, such as *Acinetobacter* and *Proteus,* which drive the high mean abundance within a given patient group (supplementary figure S1b). Overall, the abundances of these main genera were similar across patient groups (Kruskal‒Wallis test, p > 0.05).

**FIGURE 1.**
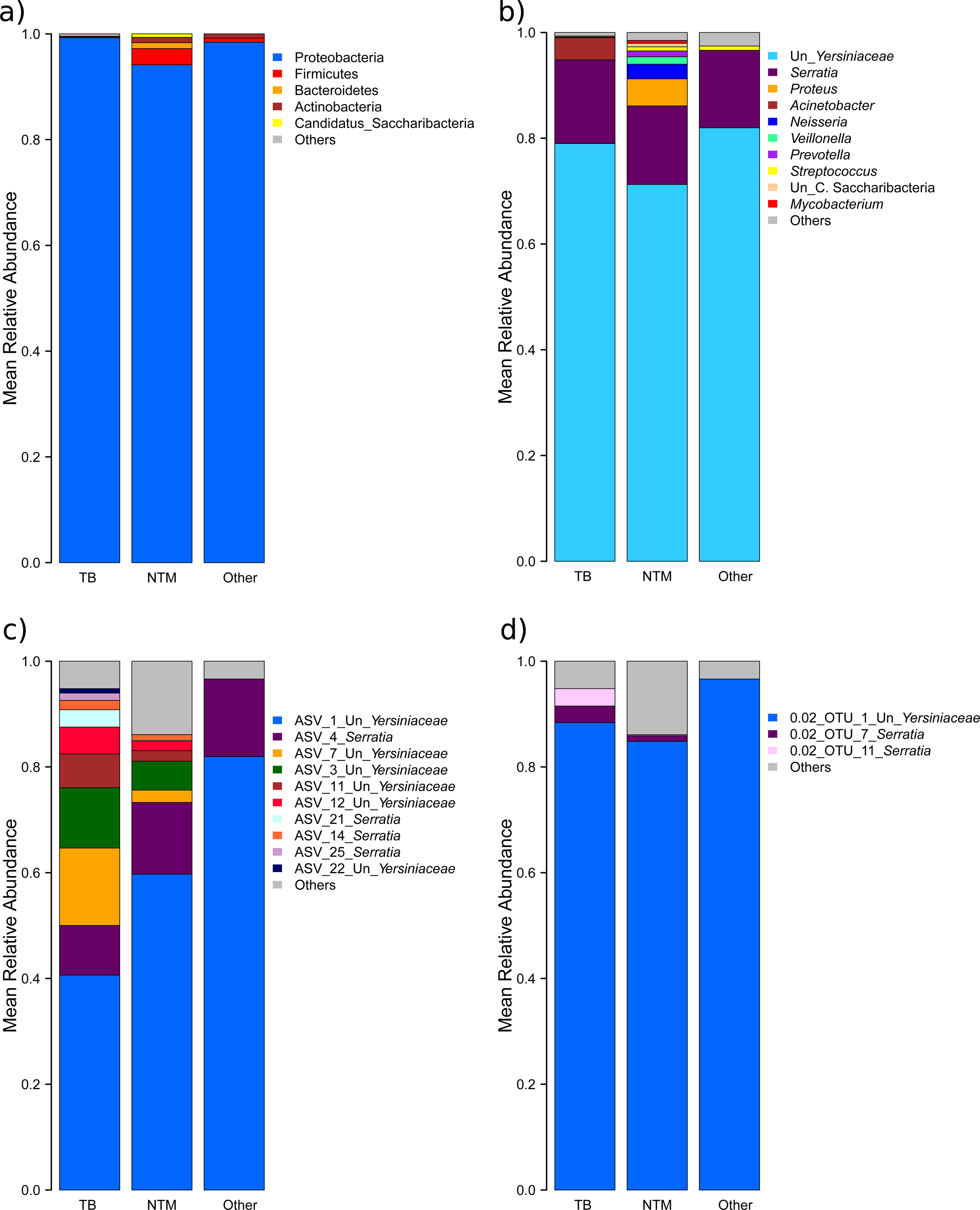
Mean relative abundances of major taxa across patient groups: a) Phyla, b) Genera, c) ASVs, d) 98% OTUs Un: Unclassified

### Diversity of Yersiniaceae and Serratia taxa

To further evaluate the diversity of the lung microbiota, and given the strong dominance of the genera **“**unclassified *Yersiniaceae”* and *Serratia* across all three patient groups, we defined taxa belonging to these genera at the sub-genus resolution level, namely the ASVs, and 98% OTUs. We compared the mean abundances of the major ASVs and 98% OTUs across the TB, NTM, patients with further pathologies. We observed that ASV_1 “unclassified *Yersiniaceae”* and ASV_4 *Serratia* dominated the communities of all three patient groups (figure 1c). Moreover, TB patients exhibited substantially greater diversity, and a significantly higher abundance of ASV_7 “unclassified *Yersiniaceae”* (Kruskal-Wallis test, corrected p= 0.042).

In addition, at the higher clustering level of 98% OTUs, we observed considerably less diversity among patients whereby a single taxon, namely, 0.02_OTU_1 “unclassified *Yersiniaceae”,* dominated the all three patient groups, while 0.02_OTU_11 *Serratia* abundances significantly differed across patients (Kruskal-Wallis test, corrected p value=0.046) (figure 1d). These results suggest that the disparities in unclassified *Yersiniaceae* and *Serratia* traits across patients lie primarily at the species/sub-species level.

To further explore the diversity of *Yersiniacea*e and *Serratia* taxa, we assessed the within-individual diversity (*i.e.* alpha diversity). Due to the substantially lower sample size (n=4) of patients with non-infectious inflammatory lung disease, we excluded this group from subsequent analyses. We found similar observed ASVs diversity between TB, and NTM-LD patients (Kruskal-Wallis test Shannon, corrected p value=0.144), and marginally significant greater expected richness of ASVs in TB-patients (Kruskal-Wallis test Chao1, corrected p value =0.0696) (figure 2a). In contrast, 98% OTUs-based diversity indices showed similar observed, and expected richness across patient groups (figure 2b) (Kruskal-Wallis test, corrected p=0.2814 for Shannon and Chao1 indices). This result is consistent with our previous observations that TB patients harbored a greater diversity of unclassified *Yersiniaceae* and *Serratia* ASVs.

**FIGURE 2.**
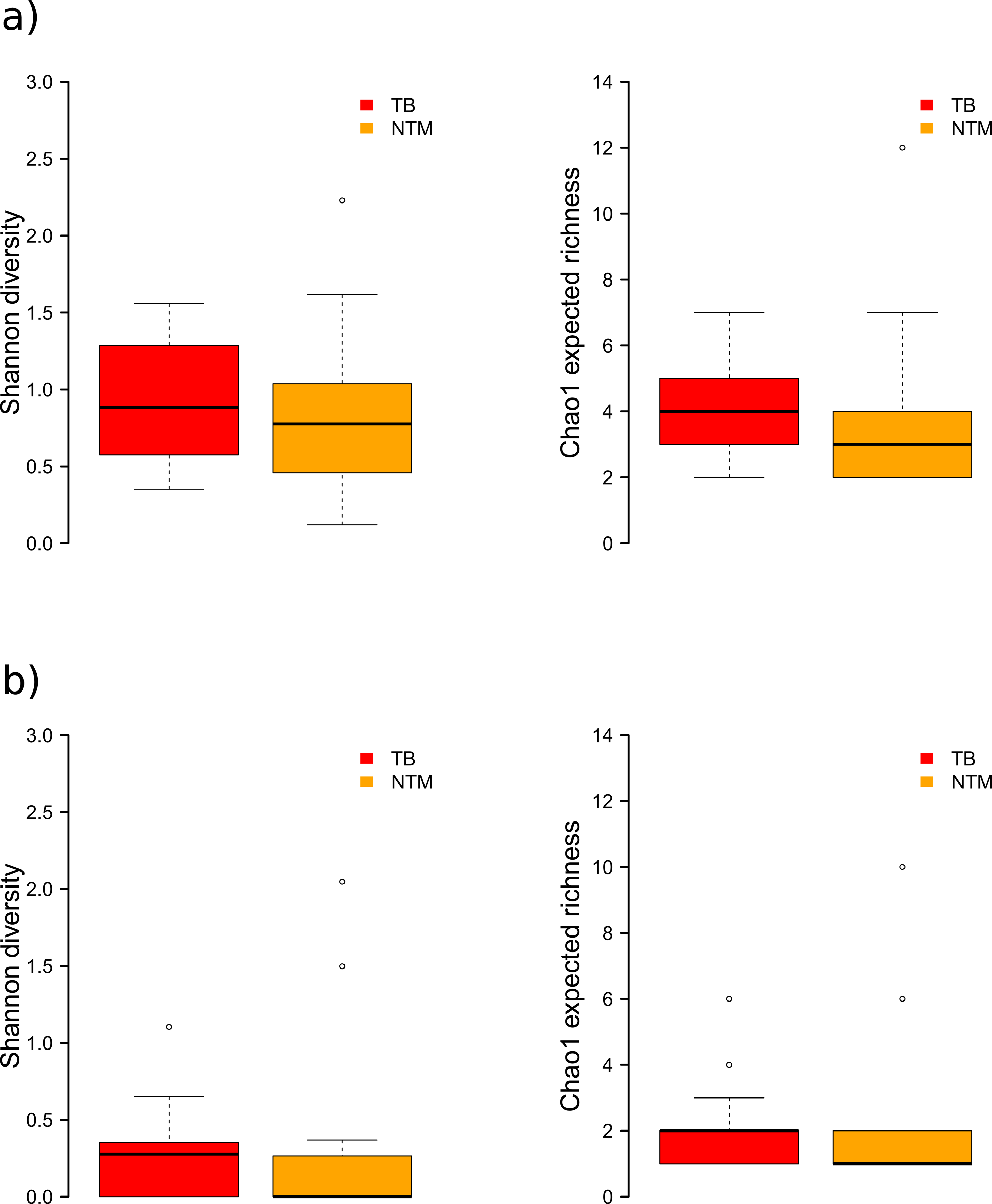
Shannon and Chao1 diversity indices in TB, and NTM patients: a) ASVs, b) 98% OTUs

### Indicator taxa of disease state

To examine potential associations between lung microbiome composition and disease, we applied “indicator species analyses” to ASVs and 98% OTUs to identify taxa which abundances differed across patient groups. First, Indicator value analyses of ASVs revealed that ASV_7 unclassified *Yersiniaceae* and ASV_21 *Serratia* tended to associate with TB-patients (corrected p= 0.17 for both ASVs). The same pattern was observed for 0.02_OTU_11 (corrected p=0.25) (supplementary table S3). Specifically, the indicator value components of specificity (A) and fidelity (B) showed that ASV_7 was highly restricted to TB patients (A= 0.86 [95% CI 0.609–1.00]) and occurred in 57% of patients (B=0.57 [95% CI: 0.360–0.767]). Similarly, ASV_21 was almost exclusively detected in BALFs of TB-patients (A= 0.94 [95% CI 0.798–1.00]) and was present in approximately 40% of the individuals (B=0.39 [95% CI 0.192– 0.60]). Furthermore, 0.02_OTU_11 *Serratia* was strongly specific to TB-patients (A= 0.94 [95% CI 0.8–0.1] with moderate fidelity B=0.39 [95% CI 0.192–0.60]).

Second, preference analysis confirmed the indicator value analyses, and revealed that ASV_7, ASV_21, and 0.02_OTU_11 significantly preferred TB-patients (p values: 0.0151, 0.039, and 0.014 for ASV_7, ASV_21, and 0.02_OTU_11, respectively, after Sidak’s correction for multiple testing). Overall, 75% the TB-patients carried one or more of the identified ASVs, and 35% harbored the indicator 0.02_OTU_1. Interestingly, these proportions changed across the specificity “A” parameter. For instance, ASV indicators, which cover 40% of TB-patients, harbored an extremely high specificity to TB-patients (A > 0.90) (figure 3a and supplementary figure S2). In summary, taxon analyses revealed distinct *Serratia* traits are exclusively identified among specimens from TB-patients. This interesting pattern likely reflect intrinsic disease and/or patient characteristics.

**FIGURE 3.**
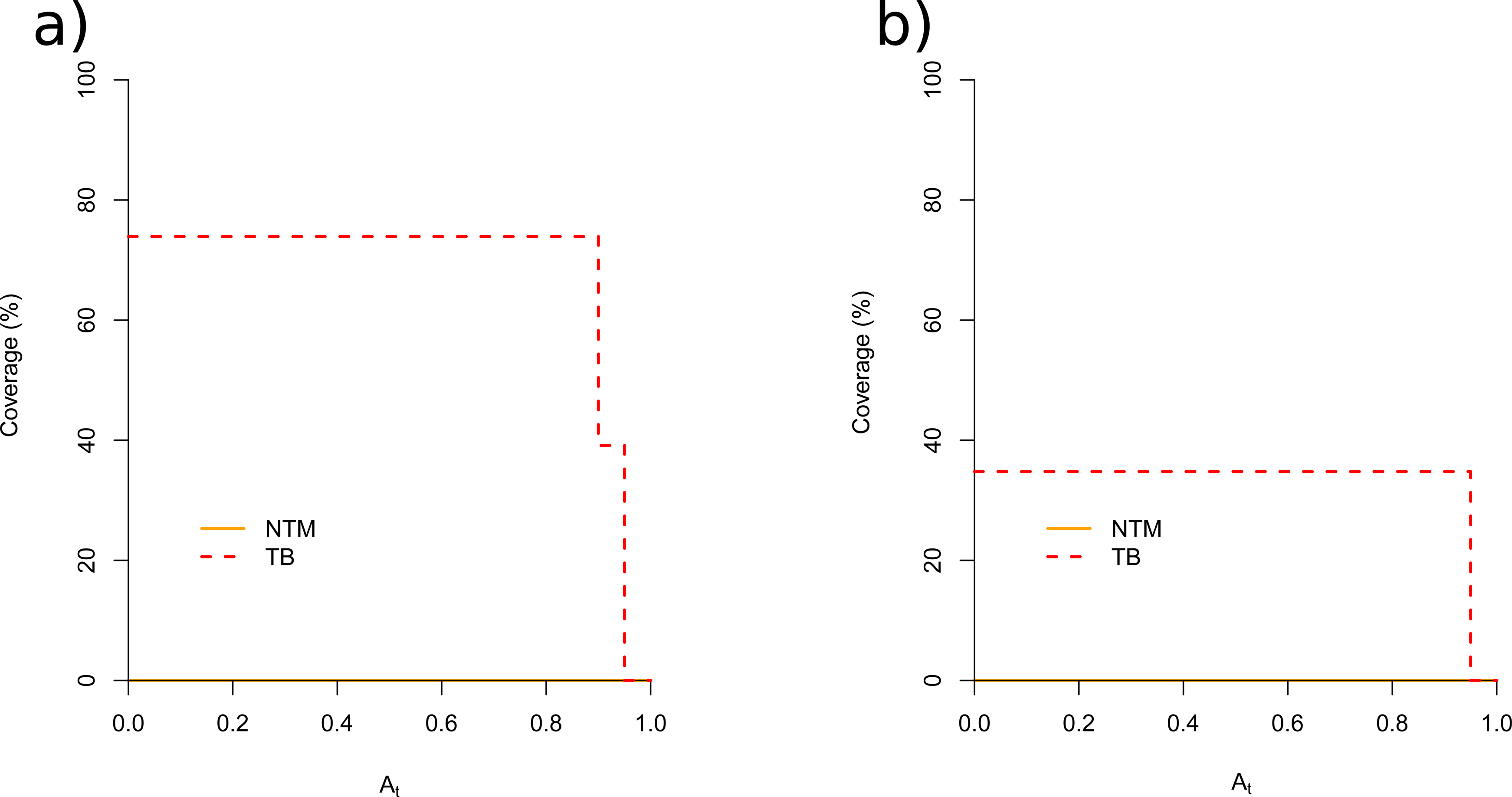
Coverage of the identified indicator taxa along the "A" threshold: a) ASVs, b) 98% OTUs

### Lung microbiome diversity across disease states

To more deeply examine the microbiome structure across disease states, we estimated Bray‒ Curtis and Jaccard indices (*i.e.* inter-individual diversity), and assessed the influence of patient, and sampling characteristics. After evaluation of the co-variates (distribution, sample size variations), only disease status, and age category were included in the final models.

In Bray‒Curtis model, disease status significantly influenced community structure, while age had a marginally significant influence (*adonis*, disease status: R^2^= 0.057, p=0.03; age category R^2=^0.219, p= 0.089, 10^5^ permutations). For the Jaccard index, disease status significantly explained a substantially greater portion of the variance compared to the Bray‒Curtis index (*adonis*, disease status R^2^ =0.072, p=0.005, 10^5^ permutations), suggesting that disparities between TB-and NTM patients were driven primarily by the presence/absence of certain ASVs. In contrast, age category did not significantly influence Jaccard index (adonis, age category R^2^ =0.166, p=0.40, 10^5^ permutations).

Furthermore, constrained principal coordinates analyses revealed that i) disease status explained a more substantial proportion of variance in the presence-absence datasets than in the abundance datasets and ii) patients clustered first by disease status, while the influence of further factors, including age category, remained marginal: Capscale, constrained inertia, Bray-Curtis=0.2403, Jaccard=0.2215; “anova.cca” by term, disease status: explained variance, Bray-Curtis=4.6%, p =0.029, Jaccard=5.30%, p=0.004; age category: explained variance: Bray-Curtis=19.76%, p=0.09, Jaccard=17%, p=0.41, 10^5^ permutations (figure 4a, and figure 4b). Of note, analyses on the 98% OTUs indicated influence of disease status remained significant only on the presence/absence datasets whereas community structure was not significantly impacted by disease status, and age category (see supplementary results, and supplementary figure S3). In essence, most disparities in lung microbiome diversity were evident at the sub-species ASV level, and disease status notably impacted community structure, particularly in presence-absence datasets.

**FIGURE 4.**
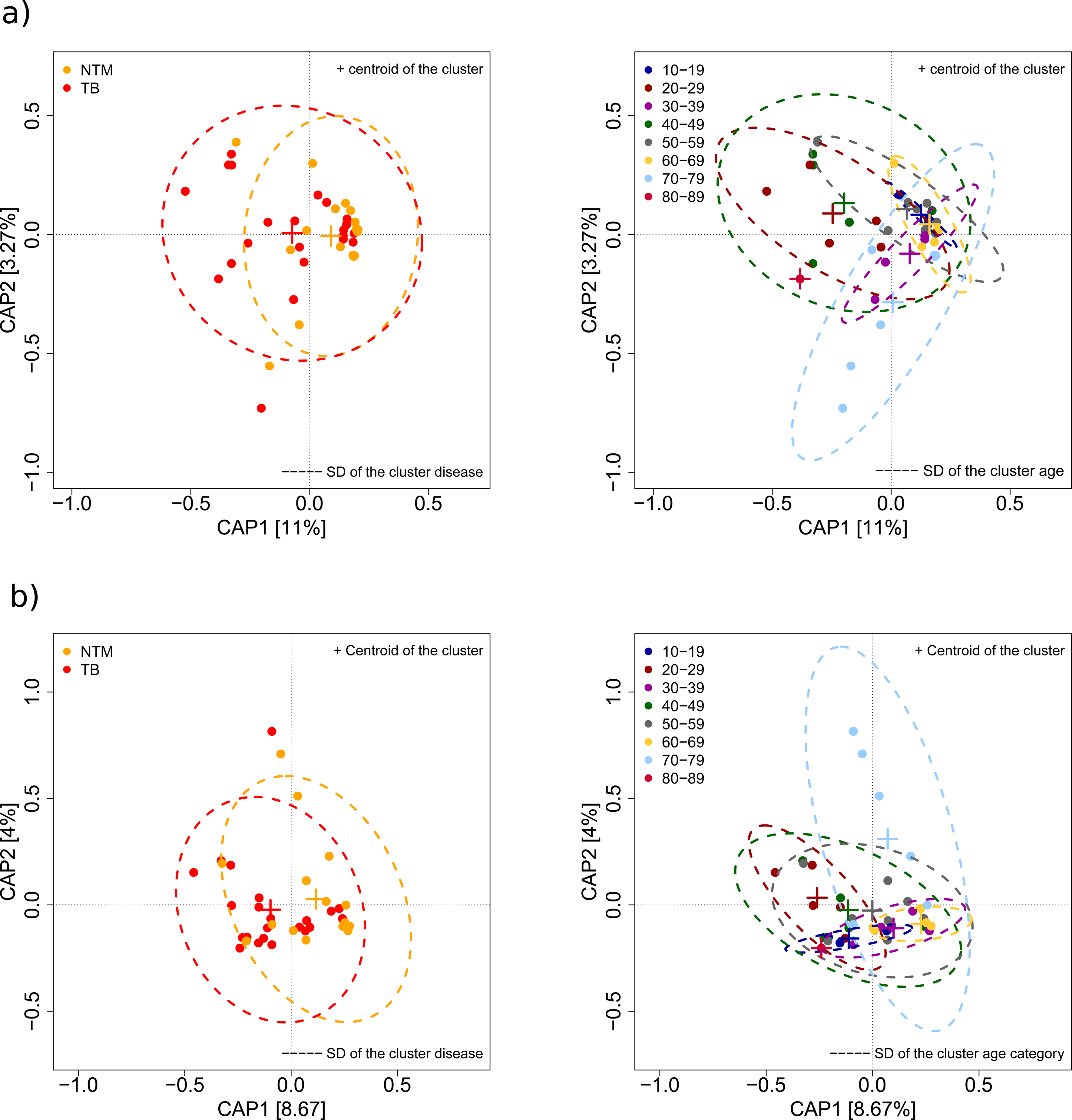
Constrained principal coordinates analysis of Bray-Curtis (a), and Jaccard (b) indices based on ASVs. SD: Standard Deviation

Lastly, we also observed distinct *Serratia* ASV and OTU interaction patterns within each disease state (see supplementary material, supplementary figures S4 and S5, and supplementary table S4). Additional exploratory analyses to determine the species taxonomy suggested that ASV_1, ASV_3, and ASV_7 represent *Serratia liquefaciens* and/or *Serratia grimesii,* and ASV_4, ASV_14, and ASV_21 indicate the presence of *Serratia myotis* and/or *Serratia quinivorans* while the WGS exploratory species assessment mainly indicated the presence of *Serratia grimesii* (see supplementary material, supplementary figure S6 and supplementary table S5).

## Discussion

We profiled the lung microbiome of patients with TB, NTM-LD, and non-infectious inflammatory lung diseases, and established a workflow for microbiome analysis of low-biomass pulmonary specimens. We confirmed previous reports showing a high abundance of the genus *Serratia* in BALF specimens. Moreover, we identified distinct *Serratia sp.* traits (*e.g.*, subspecies or strains) that characterized and distinguished the lung microbiome of patients with pulmonary TB, and NTM-LDs. However, it remains to be characterized whether individual strains of *Serratia sp*. and pathogenic mycobacteria interact mechanistically, and whether their co-occurrence in the lower airways influences the onset or progression of disease.

### *Yersiniaceae* and *Serratia* dominate the lung microbiome of pulmonary patients

We revealed that the genera unclassified *Yersiniaceae* and *Serratia* (family *Yersiniaceae*) dominated the lung microbiome of individuals with pulmonary TB-patients, NTM-LD, and non-infectious inflammatory lung diseases. Similarly, previous studies that combined BALF and 16S rRNA amplicon sequencing reported a high prevalence of specific single taxa [20, 21]. Specifically, *Serratia* has been described in diverse lung pathologies. Gupta and colleagues [22] reported that *Serratia* dominated the lungs of patients with exacerbated COPD, interstitial lung diseases and sarcoidosis but not in patients with stable COPD. In a large cohort of individuals with pulmonary TB, Hu et al. [23] identified the genus *Serratia* as the major taxon in the lung microbiome of patients with or without culturable mycobacteria from BALFs. Differences in the microbiome composition before, and after anti-TB therapy, *i.e.*, after culture conversion, were partially driven by two *Serratia* OTUs. The OTUs abundances decreased after sputum culture conversion, indicating that *Serratia* traits were affected by the presence/absence of Mtbc bacteria and/or the anti-TB medication itself. Notably, Hu et al. [26] and others could not detect the genus *Mycobacterium* in all culture-positive specimens, highlighting the detection limits of 16S rRNA sequencing for these low-biomass specimens [24, 25]

### TB-patients harbor a distinguishable heterogeneous lung microbiome structure

Our data confirmed previous studies, using 16S rRNA amplicon sequencing and BALFs or lung biopsy, that the lung microbiome of TB and NTM-LD patients prior to therapy exhibited greater richness as compared to controls [26–28]. In contrast, Xiao et al. [29] employed WMS and reported increased diversity, and expected richness in cured TB-patients, while patients receiving therapy exhibited intermediate patterns, and active TB-patients showed the lowest diversity, and richness. Overall, the observed higher richness in (untreated) TB-patients is likely the consequence of impaired colonization resistance, whereby the healthy homeostatic microbiome prevents the infiltration, and growth of opportunistic pathogens [30–32]. The start of anti-TB therapy profoundly alters the lung microbiome structure, further decreases potentially beneficial taxa, and selects for antibiotic resistance genes (ARGs) [28, 29].

To date, it is unclear whether certain strains or subspecies can exploit this niche, and possibly affect lung pathology, disease progression or treatment outcomes. In addition to the elevated taxa richness in TB, our analyses of beta diversity measures confirmed that TB, and NTM-LD significantly influenced community structure prior to therapy, and more strongly on presence/absence, while the impact of patient-related factors, including age, remained marginal. Moreover, we identified specific ASV signatures for the lung microbiome of TB-patients, however, we also highlight the individual microbiome heterogeneity especially of TB-patients.

### *Yersiniaceae* and *Serratia* harbor distinctive traits, with different interaction patterns

Our data further suggested that more than one strain or subspecies of *Serratia sp.* was present in the lung microbiome of patients. To date, the mechanisms underlying intraspecies and interspecies interactions within the genus *Serratia* remain poorly understood, and are largely focused on the opportunistic pathogen *Serratia marcescens* [33–35]. Strikingly, pioneer studies characterized the molecular basis of interactions between *S. marcescens* and *M. tuberculosis* and demonstrated a growth-promoting effect of specific siderophores secreted by

*Serratia marcescens* in iron-depleted environments. Further cytotoxic assays revealed that these siderophores exhibit cytotoxic activity against human cells [36, 37]. Here, we observed that within patients with pulmonary TB and NTM-LD, *Yersiniaceae*/*Serratia* ASVs exhibited differential interaction patterns, with correlations shared across TB and NTM-LD patients, whereas others were unique to a specific disease state. This intriguing pattern likely indicate a dynamic disease-specific interaction driven by growth competition among distinct subspecies and/or strains of *Yersiniaceae*/*Serratia*, as demonstrated for other barrier organs, including the gut and skin [38–40]

### Strengths and limitations

We reveal the disparities, and heterogeneity of the lung microbiome of TB-compared to NTM-LD patients, and present *Serratia sp.* traits as a major player in understanding the microbiome dynamics in Mtbc, and NTM lung infections.

Due to the retrospective study design, we could not collect samples from negative controls such as saline oral and bronchoscope washes. Thus, we constructed a rigorous and strict decontamination scheme. Specifically, we assessed the bacterial biomass in the BALF supernatants to detect possible contaminants, *i.e.*, taxa that negatively correlated with the sample biomass and were not detected across technical replicates. In addition, we included several positive controls (mixed and pure cultures), and technical replicates of positive controls and native BALFs, to optimize the detection of contaminants and spurious taxa, as recommended earlier [13].

### Conclusion and outlook

*Serratia* has emerged as a potential crucial player in understanding the dynamics of the lung microbiome in pulmonary TB and NTM-LD. Further work is needed to explore strain-level effects among the genus *Serratia*, microbe‒microbe interactions, and resulting implications for disease progression and therapeutic outcomes.

## Acknowledgments

We thank Anja Lüdemann, Larissa Mohr, Vanessa Mohr, Maja Mundzeck, Tanja Niemann, Silvia Maass, and Franziska Daduna for excellent technical assistance.

## Ethics statement

The study was positively evaluated by the ethics committee of the University of Lübeck (EK HL AZ 22-249).

## Data availability

The raw 16S rRNA amplicon-, and the short-read sequences were submitted to the Sequence Read Archive (SRA) under BioProject PRJNA1103672

## Author contributions

**MB** designed the study, optimized the laboratory protocols, designed analysis strategy, performed the statistical analyses, and co-wrote the paper. **BK** recruited the patients, performed the bronchoscopy, and collected the BALF specimens. **SN** contributed to discussion and provided WGS infrastructure. **KIG** provided patients metadata from the biobank, and the ethics declaration. **CL** received the patients, performed the bronchoscopy, and collected the BALF specimens. **JH** designed the study, received the patients, performed the bronchoscopy, and collected the BALF specimens. **MM** designed the study, performed the WMS analyses, and co-wrote the manuscript

---

**S. FIGURE 1.**
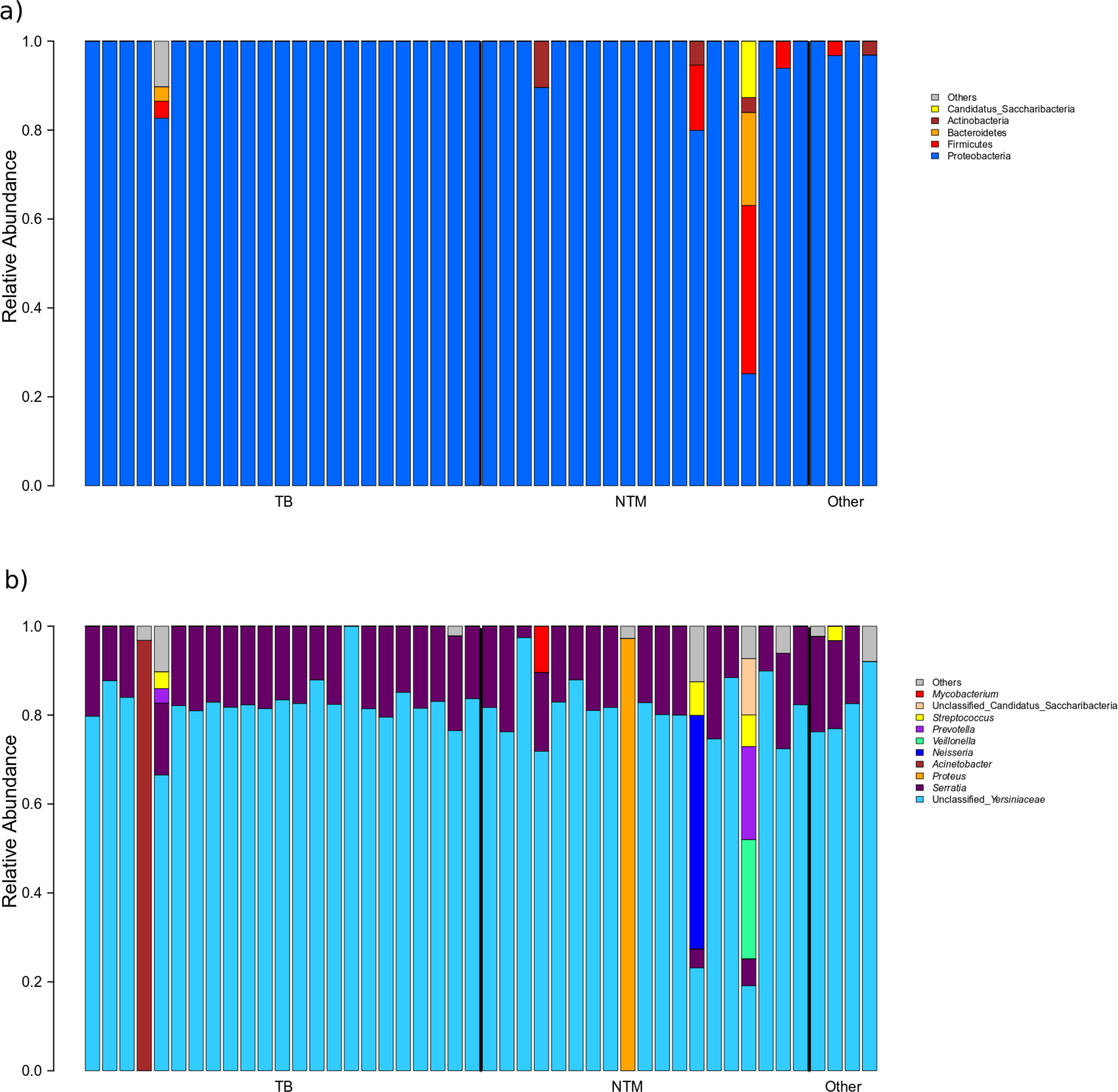
Relative abunda1nces of major taxa: a) phyla, b) genera across patient groups

**S. FIGURE 2.**
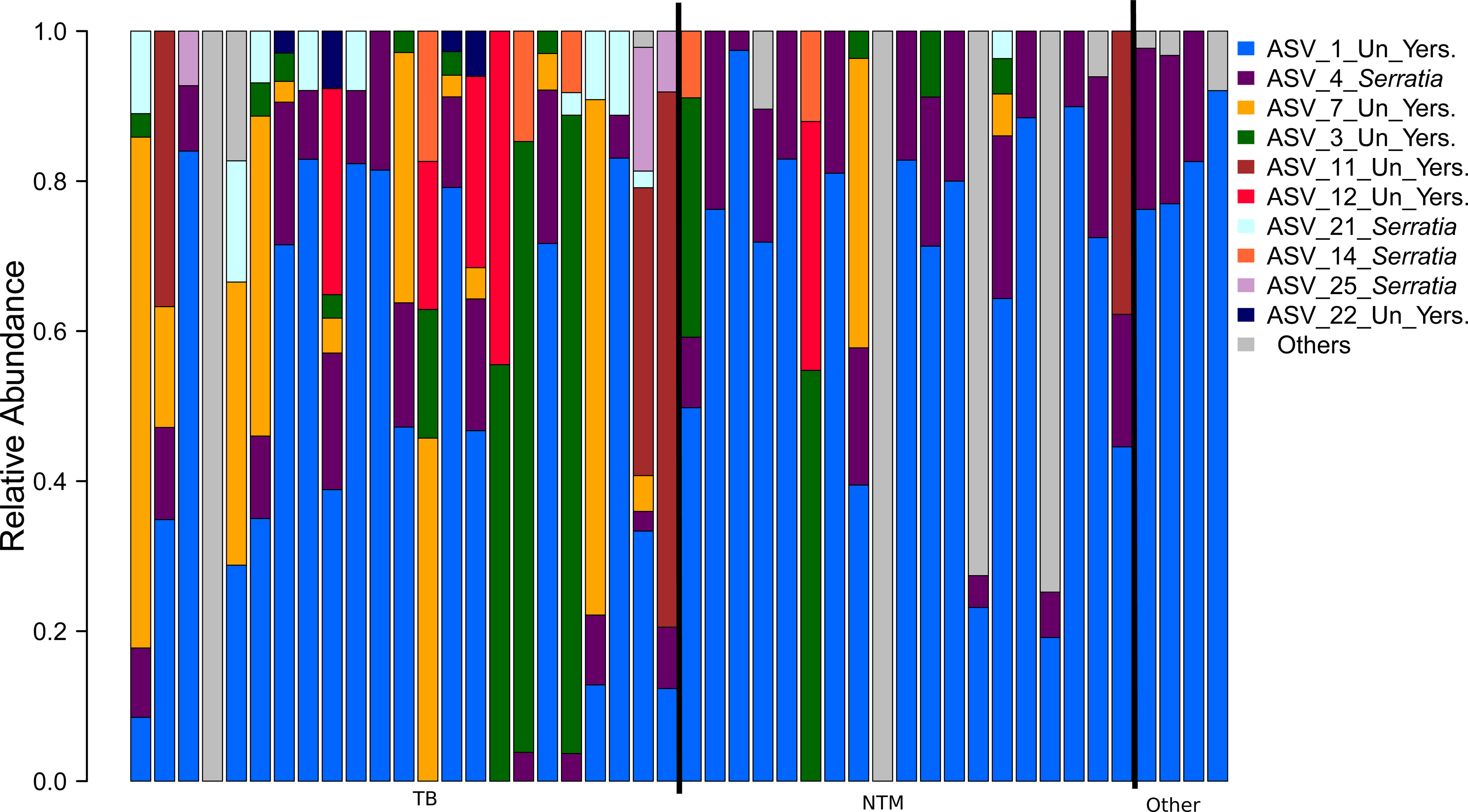
Relative abundances of unclassified *Yersiniaceae*, and *Serratia* ASVs. Un: Unclassied. Yers: *Yersiniaceae*

**S. FIGURE 3.**
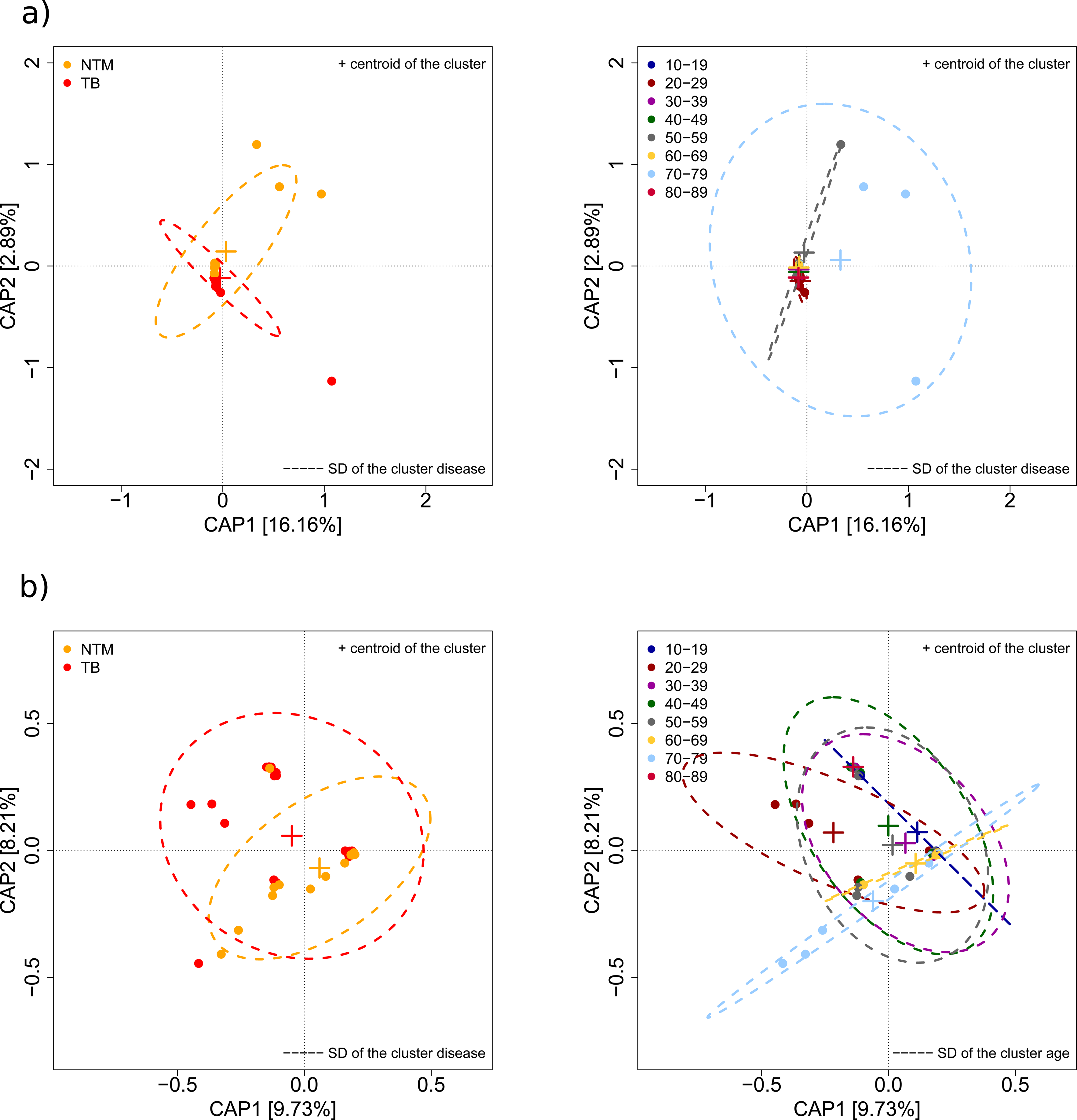
Constrained principal coordinates analysis of Bray-Curtis (a), and Jaccard (b) based on 98% OTUs

**S. FIGURE 4.**
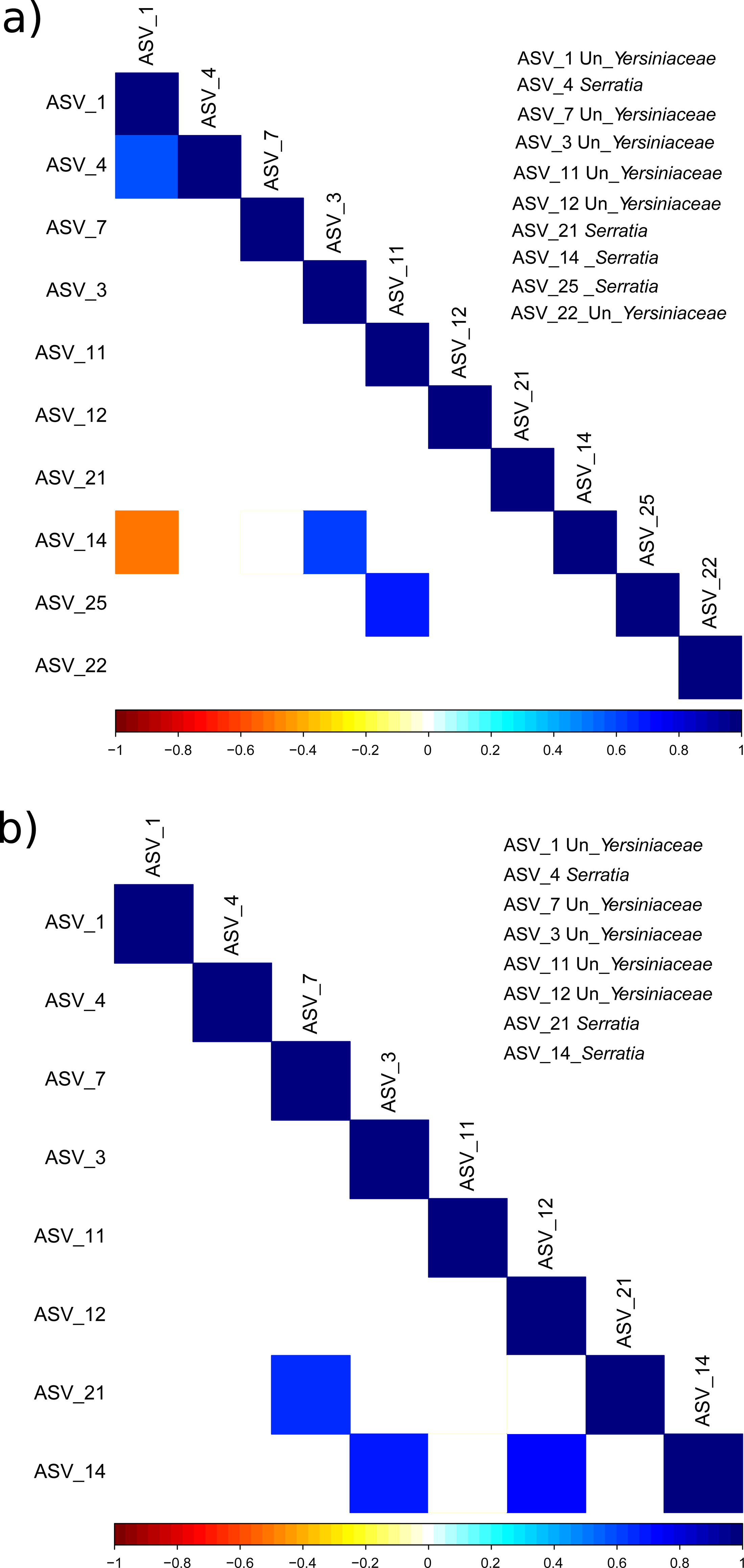
Pairwise Spearman’s correlation matrix between major ASVs: a) TB, b) NTM Only significant correlations after p values correction are shown. Un: Unclassified

**S. FIGURE 5.**
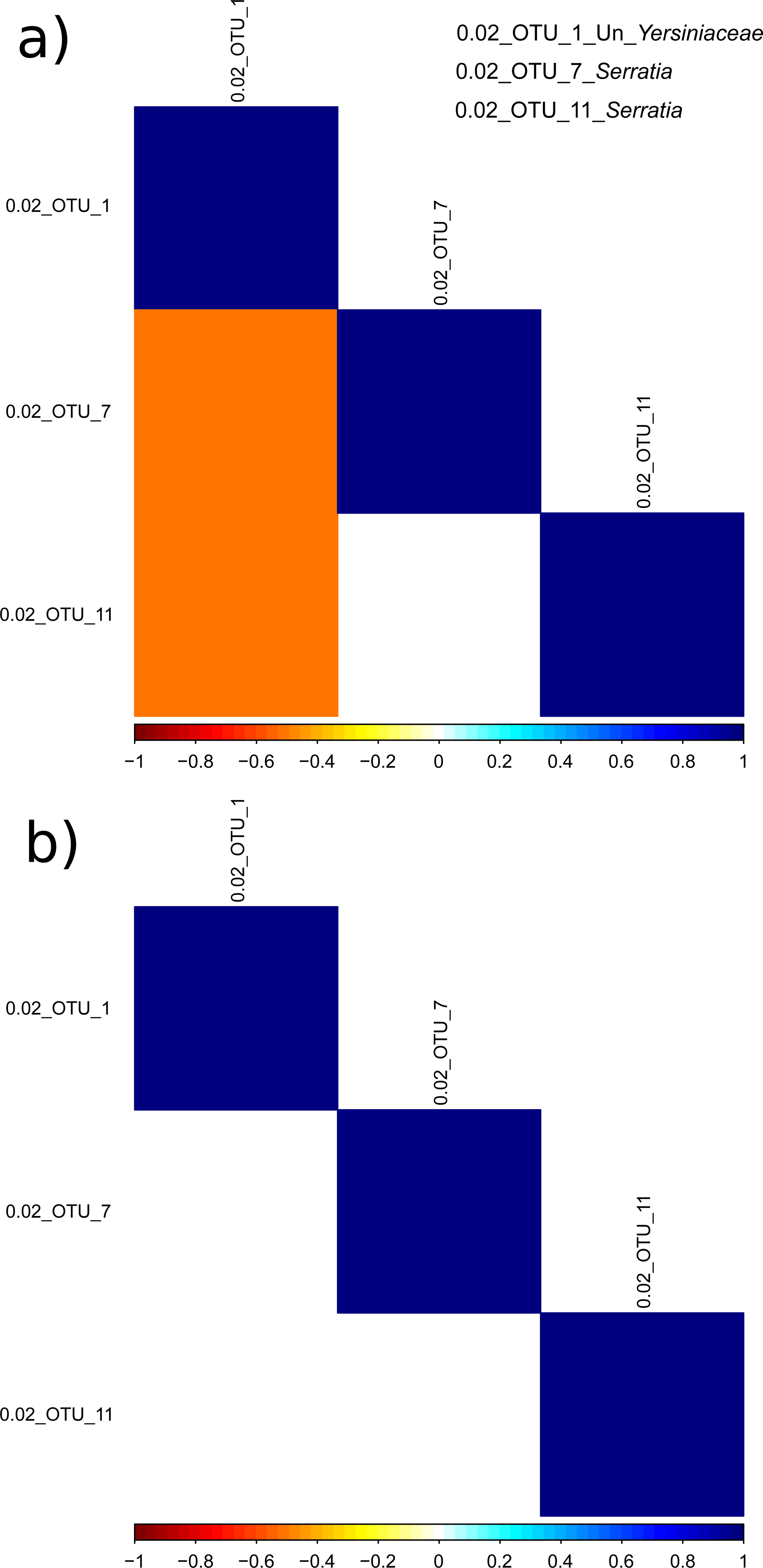
Pairwise Spearma’s correlation matrix between major 98% OTUs: a) TB, b) NTM. Only significant correlations after p values correction are shown . Un: Unclassified.

**S. FIGURE 6.**
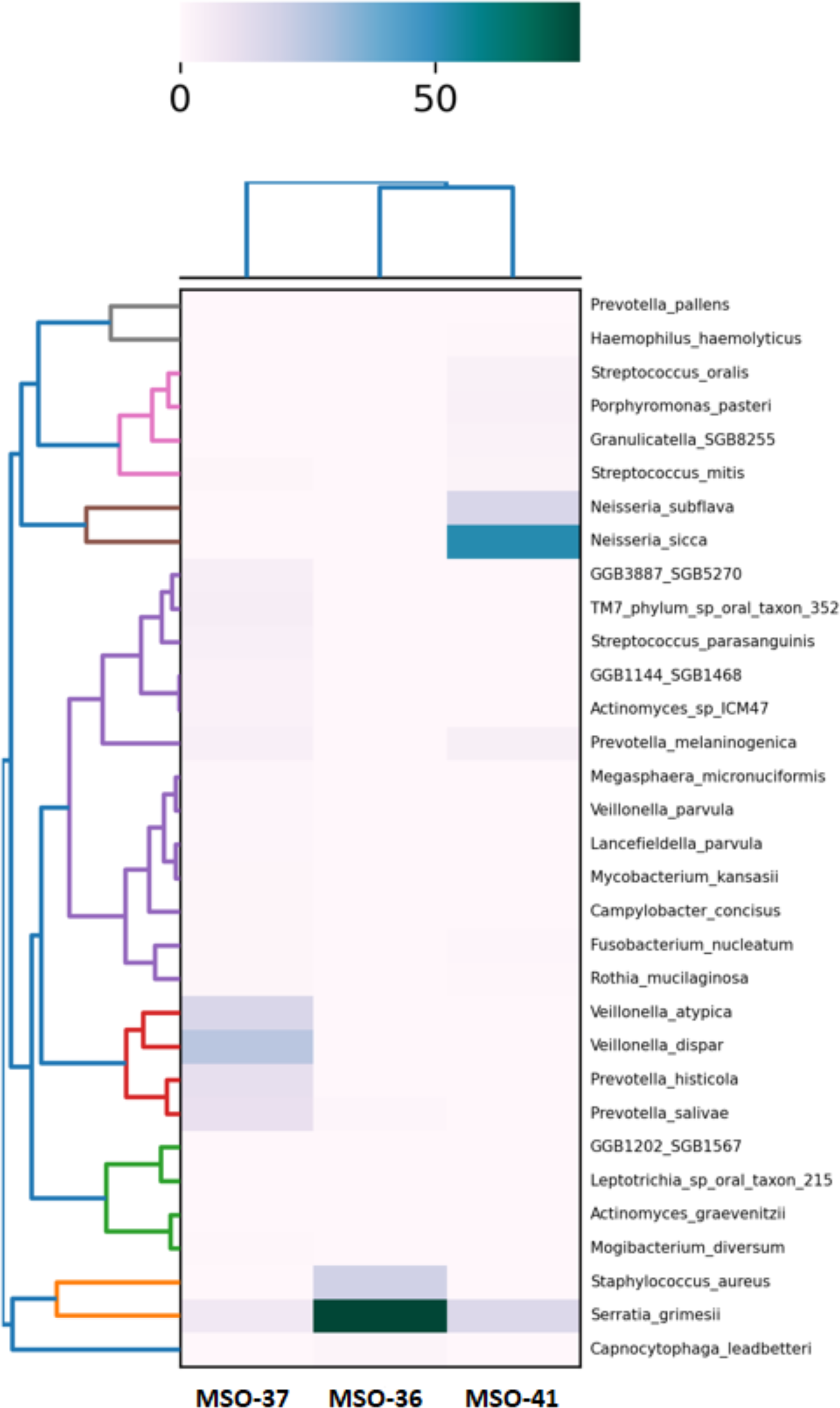
Relative abundances of microbial species identified in 3 NTM patients. Only species with a relative abundance >0.1% are shown, with *Serratia grimesii* occuring in all 3 BALF specimens (from left to right: 7%, 78%, and 16%, respectively)

