## Supplementary Material for "*Serratia sp*. dominates the lung microbiome of patients with tuberculosis and non-tuberculous mycobacterial lung diseases"

**Supplementary Methods**

**Depletion of host cells, extracellular DNA, and DNA extraction**

The total BALF was first subjected to a differential centrifugation step to facilitate the separation of host and microbial cells at 1800 rpm for 10 minutes at 4°C. Approximately 10-50 ml of the supernatant was collected and stored at -80°C until DNA extraction. Subsequently, the acellular BALF supernatants were first thawed at 4 C° overnight, then centrifuged at 5000 rpm to collect the bacterial cells, at 4 C° for 20 minutes. After removal of the supernatant cell pellets were resuspended in 500 µl PBS. An equal volume of PCR-grade water was added to the re-suspended cells pellet, gently mixed, and incubated at room temperature for 5 min to osmotically lyse mammalian cells (Marotz et al, 2021). Thereafter, 1 µl Turbo Dnase (2 units) mix (TURBO™ DNase (2 U/µl), Invitrogen) was added to the BALF cells pellets, and incubated at 37 C° for 30 minutes, followed by centrifugation at 10,000xg for 10 minutes at 4 C°. After removing up to 400 µl supernatant, cells pellet was transferred to a PowerBead tube (DNeasy UltraClean Microbial Kit, Qiagen) containing 300 µl bead – beating buffer, 50 µl SL buffer, and 20 µl proteinase K (20mg/ml). PowerBead tubes were subjected to homogenization in a the FastPrep-24 instrument at 6.5 M/S speed for 15 seconds, three times. Subsequent DNA extraction steps were conducted according to the manufacturer’s protocol, and DNA was eluted in 50 µl EB solution. Blank extraction negative controls were included in each extraction round. In addition, a sequence of an 8-fold dilution series (named from D0 to D8) of the ZymoBIOMICS microbial cells standard (D6300) was extracted along the BALFs to assess the contamination thresholds along microbial DNA concentrations as recommended by Karstens and colleagues [1].

**Further positive controls, and technical replicates**

Beside BALFs, blank extraction controls, and dilution series of the microbial cells’ standard, we included various types of positive controls: i) isolates of pure cultures previously sequenced, and characterized: *Haemophilus haemolyticus*-DSM-103601, *Haemophilus influenzae*-DSM-11121, *Mycobacterium abscessus*-21002932-ATCC19977, *Mycobacterium abscessus*-2100299, *Mycobacterium abscessus*-21003041, *Mycobacterium*-22003835, *Mycobacterium*-22003836, *Mycobacterium*-Positive-C2, BCG-Positive-C1, ii) the microbial community DNA standard (10 ng/μL) (ZymoBIOMICS). Furthermore, we generated technical replicates (same biological sample amplified, and sequenced several times, independently) as an additional tool to detect contaminants in BALFs samples [2]. In total 16 different samples (pure cultures n= 4, BALFs n= 11, and Mock standard n=1) were replicated 2, 3, 5, or 6 times.

**16S rRNA amplicon sequencing, and processing**

The V3-V4 hypervariable region of the 16S rRNA gene (around 465 bp) was amplified following a dual-indexing approach using barcoded 341F and 806R primers carrying unique eight-based MIDs, and heterogeneity spacers [3]. The 25 µl-PCR reaction included 9.25 µl PCR-grade water, 0.5 µl dNTPs mix (10µmol), 0.25 µl Phusion™ High-Fidelity DNA Polymerase (2 U/µL), 5 µl Buffer (5X), and 2 µl DNA. 4 µl of each forward, and reverse primer (2 µmol). Thermocycler cycling conditions were as follows: initial denaturation 98 C° for 5 minutes, 98 C° for 9 seconds, 55 C° for 60 seconds, 72 C° for 90 seconds, 30 cycles; final elongation at 72 C° for 10 minutes. PCR negative controls were included for each forward, and reverse primer combination, checked for amplification on a 5200-fragment analyzer system (Agilent) using the DNF-930-Kit, and required to be free of amplification: if an amplification was detected, the PCR reactions of all samples amplified using those primers are repeated with freshly prepared reagents. PCR-products were cleaned up the using MagSi-NGSPREP Plus (magtivio) Left Side Size Selection protocol, then checked on the 5200-fragment analyzer system (Agilent). Finally, Purified PCR-products were quantified using the Qubit 1X dsDNA HS Assay-Kit (ThermoFisher Scientific), on a Qubit Flex Fluorometer, and mixed in a final pool at an equal molar concentration (2 nM). The pool library was further diluted to 12 pm, denaturated, and sequenced along with 50% PhiX on the illumina Miseq platform using the V3 Miseq sequencing kit 2 X 300 cycles. Negative extraction controls did not yield amplification and were not sequenced.

Raw Fastq files were processed in dada2 R package (v.1.16.0) [4] as follows: forward, and reverse reads were truncated at 270 bp, and 250 bp length, respectively. After truncation, reads with any ambiguity, and maximum number of expected errors maxEE higher than 3 were discarded. Trimmed forward, and reverse reads were first dereplicated, denoised, and merged with parameter minOverlap = 20, then the Amplicon Sequence Variant (ASV) table was defined. ASV Chimeras were identified, and removed within each sample independently using method “per-sample”. Taxonomy was assigned to each ASV from phylum to genus with minimum bootstrap confidence minBoot = 80 using the ribosomoal database project (RDP) classifier (release 18) [5]. Finally, reads classified as non-bacteria, and reads inaccurately merged (shorter than 418 bp, and longer than 512 bp) were excluded. This yielded to 12987 defined ASVs across BALFs, negative, and positive controls.

**Adjusting ASVs clustering**

Although ASVs more accurately characterize contaminants in low biomass samples [6], clustering sequences into ASVs might split genomes, and thus inflate the diversity of microbial types detected within samples [7]. Thus, to correct for potential misidentified ASVs, we inspected the pure cultures, the Mock standards, as well as the technical replicates to evaluate the accuracy of the ASVs in quantifying the composition, and expected diversity within samples. We observed that i) across the pure cultures (n=21), 23 ASVs were defined on average within a single sample, ii) the Mock samples (n=8) (expected 8 different species) showed on average of 150 ASVs, iii) the technical replicates (n=16) showed poor to no overlapping ASVs (Supplementary Table 1). Therefore, to correct the sequences clustering to better reflect the samples true diversity, we re-clustered the originally defined ASVs to corresponding adjusted ASV following this thorough procedure:

- **Step 1**: **Alignment of ASVs sequences, and calculation of a pairwise distance matrix**

All defined ASV sequences (n= 13054) were aligned in R package DECIPHER (2.24.0) [8] using function “AlignSeqs” with 10000 interactions, and 1000 refinements. Subsequently, a pairwise distance matrix was calculated based on the alignment using “DistanceMatrix” function with includeTerminalGaps=TRUE.

- **Step 2: Define the range of ASVs pairwise-distances detected within pure culture**

On the subset of pure cultures (n=21) (9 different samples, and their technical replicates assessed, distinctively): First, we collected pairwise distances between the ASVs detected within each sample, and which classified to the expected genus (to exclude potential contaminants). Second, we defined the summary statistics calculated on all collected pairwise distances collected across the samples, including minimum, first quartile, median, mean, 3^rd^ quartile, maximum, and standard deviation.

- **Step 3: Define the best-fitted distance to cluster ASVs into one adjusted ASV**

To define the optimum pairwise distance at which ASVs should be clustered to one adjusted ASV, and better reflect the true composition, and diversity within samples, we independently tested the distances defined in step 2, namely median, mean, 3^rd^ quantile, 3^rd^ quantile + standard deviation, and the maximum pairwise distances. Precisely, ASVs which pairwise distance equaled or were inferior than the tested distance, were grouped, and assigned to the same adjusted-ASV, and the most abundant ASV was defined as the lead ASV of the newly defined adjusted-ASV cluster. Accordingly, and following a sample-wise manner, abundances table of the adjusted-ASVs across samples were defined by adding up the abundances of the ASVs assigned to a given adjusted-ASV.

- **Step 4: Select the best-fitted distance to cluster the ASVs into one adjusted ASV**

To select the optimum clustering pairwise distance among the tested distances in step 3, we examined the ratios of shared, and non-shared/unique adjusted-ASVs within each pure culture sample (n=4), and across its technical replicates (between 2 to 6 / pure culture sample), for each of the tested clustering pairwise distances. We found that the maximum pairwise distance (0.0168) yielded the highest fraction of shared adjusted ASVs across the technical replicates of a given pure culture. Afterwards, we further checked, and validated the accuracy of the adjusted ASV clusters based on the maximum pairwise distance: i) All ASVs grouped within the new adjusted ASV had identical taxonomy, ii) the fraction of shared adjusted ASVs within the BALF samples, and their replicates drastically increased compared to the shared ASVs, iii) the frequency/number and taxonomy of the adjusted ASVs in the Mock standards was closer to the expected composition, and diversity (Supplementary Table 1). This final ASVs clustering resulted in 2084 adjusted ASVs in the entire dataset, for all subsequent analyses.

**Detection of contaminant, and spurious ASVs**

- **Reference of true resident, and contaminant ASVs**

With the aim to detect contaminants, spurious sequences, and define the minimum amplified microbial DNA sufficient to reliably decontaminate our experimental samples, we defined a reference table of true residents, and contaminants based on the Mock standards dilutions as follows: We retrieved the 16S rRNA gene sequences of the species present in the Mock standard, ii) aligned sequences of all adjusted ASVs detected (n=124) across the mock dilutions (n=7) D0x2, D1, D2, D3, D4, and D8 with the retrieved expected 16S rRNA sequences in Geneious Prime (v. 2023.0.4) [9] using the Clustaw algorithm (Clustal Omega 1.2.2, Number of refinement iterations = 100, Cluster size for mBed Trees n= 100).

Next, aligned sequences were trimmed to equal lengths, and a distance matrix was calculated based. Each of 124 Mock adjusted ASVs was assigned a status of “Contaminant” or “True resident”: if an ASV sequence had an equal or smaller than 0.03 pairwise distance to a Mock standard retrieved sequence, it was assigned “true resident” status. On the contrary, if an ASV sequence showed larger than 0.03 pairwise distance to all Mock retrieved sequence, it was assigned as “Contaminant”. Overall, 23 ASVs were defined as true residents, 95 contaminants, and 6 ambiguous on the 124 adjusted ASVs detected across the Mock standard dilutions.

- **Identification of contaminant ASVs using “decontam”**

We used “decontam” (v.1.16.0) [10] in R package phyloseq (v.1.40.0) [11] to identify contaminant sequences with the frequency method which null hypothesis is “sequence A is a contaminant”. We used the FA fluorescent intensities of the purified amplicons measured prior to combining the samples into an equimolar pool as a proxy for DNA quantity. We applied this method on adjusted ASVs dataset (all samples, n=2084 adjusted ASVs), and tested several probability thresholds 0.1, 0.2, 0.3, 0.4, and 0.5. We assessed the accuracy of each threshold using the Mock standard dilutions, and the reference table of true resident and contaminants defined in the previous step. The optimum threshold was selected based on: i) ratio of false negatives (identified as contaminant, but is a true resident member), and ii) false positives (identified as a resident member, but is a true contaminant). Threshold 0.4 detected most contaminants while not defining false negatives (identified as contaminant, but is a true resident member), and thus was selected. We detected 206 contaminants adjusted ASVs.

- **Minimum abundance threshold of adjusted ASVs**

Since decomtam algorithm is not sufficient to detect all contaminants, and likely not the cross-contaminants which are typically present at low abundances [1], and abundance thresholds of spurious sequences should be study- and samples type specific [12]; to set the minimum abundance at which an ASV is reliably detected within a sample, we used the technical replicates (n=16: 4 pure cultures, 11 BALFs, and 1 mock D0), and evaluated the abundances of adjusted ASVs which replicated/shared within a sample, and between its technical replicates. We found that the mean relative abundances of shared adjusted ASVs varied greatly across sample types: 50% for pure cultures, and 6% for BALFs. Thus, we tested median, mean, and 3^rd^ quantile abundances for the shared adjusted ASVs as defined based on the BALF replicates, besides thresholds 1%, and 2%.

Following a sample-wise manner, we excluded (set abundance to 0) each adjusted ASV which relative abundance was lower than the tested abundance threshold. Next, we determined the best-fitted minimum abundance threshold based on the Mock standard dilutions reference previously defined, and found that threshold 2% showed the best ratio of removed true contaminants, and retained true residents. Thus, the 2% minimum abundance was applied sample-wise on the entire dataset: if a within a given sample, an adjusted ASV had a relative abundance lower than 2%, its abundance was set to zero within that sample. Overall, 1898 adjusted ASVs were identified as spurious, meaning they were present across all samples at abundances lower that 2%. Finally, we combined the defined contaminants (using decontam), and spurious (using the minimum threshold method) adjusted ASVs (n=1945), and excluded them from the entire dataset. This resulted in 139 decontaminated reliable adjusted ASVs.

- **Efficiency of the decontamination, and minimum DNA quantity**

To define the minimum DNA quantity at which a sample is reliably decontaminated, we examined the contamination percentages across the mock dilution series, and found an average of 3.61%, a minimum of 0.29%, and a maximum of 7.73% for dilution D8 which was efficiently decontaminated (all remaining detected ASVs across all dilutions were true residents) as described above. Accordingly, all BALF samples (n=9) which DNA concentration was lower than the D8 concentration were excluded. This, and after excluding BALF samples, n=123 adjusted ASVs defined in the dataset.

**Normalization of sequencing depth, and 98% OTUs clustering**

To normalize the sequencing depth across samples, and reduce biases in subsequent/following ecological estimates/analyses, we randomly 10 000 reads from each sample. The corresponding normalized adjusted ASVs, genera, and phyla abundance tables were adapted/modified accordingly. Samples with lower than 10 000 reads were excluded, and final analyzed samples size was n= 46 unique BALF sample, NTM =19, TB= 23, and 4 others. Subsequently, to explore the microbiome composition at a sub-genus, yet above-ASV level, we further clustered the adjusted ASVs into 98% OTUs as described above whereby sequences which pairwise distance equaled or was less than 0.02 were assigned to the same 98% OTU cluster. Within each 98% OTU cluster, the most abundant adjusted ASV was defined as the lead ASV. The same procedure was repeated with pairwise distance equaled or was less than 0.03 to define 97% OTUs.

**Ecological and statistical analyses**

The mean relative abundances of the main phyla, genera, adjusted ASVs, and 98% OTUs were compared across patient groups using the nonparametric Kruskal‒Wallis test. Correction for multiple testing was performed using Benjamini and Hochberg’s method (Benjamini and Hochberg, 1995) [13]

To identify indicator taxa of disease states, we applied indicator value analysis using the function “multipatt” in the “indicspecies” R package (v.1.7.13) [14] with func = “IndVal.g” and 10^5^ permutations. The confidence intervals of the indicator value components, *i.e.* the strength of taxon-patient group associations, were evaluated using the function “strassoc”, with alpha.ci = 0.05 and nboot.ci= 10^5^. Moreover, we applied the “signassoc” function to assess the preference of a taxon for a target group, independently from the strength of the association value (H0, the preference of the species for a given site group is due to chance only), with mode=1, two-sided alternative hypothesis, and 10^5^ permutations. Besides, we evaluated the proportion of individuals in the target patient group in which one or another indicator was found and applied the function “coverage”, with type=”stat”, At=0.5, B=0.2, alpha = 0.05, then examined the extent to which individuals in a given patient group are covered by the identified indicators along the “A” threshold using the function “plotcoverage”.

To explore the interactions between different taxa within patient groups, we calculated Spearman´s correlation between the relative ASV and 98% OTU abundances. We used “Hmisc” (v.5.1.0) [15] to calculate correlation coefficients, and “corrplot” (v.0.92) [16] to visualize significant (p ≤ 0.05) correlations. P values were adjusted for multiple testing as described by Benjamini and Hochberg [13].

**Whole-metagenome sequencing (WGS)**

DNA libraries with an average size of 700 bp were prepared as described earlier [17] (Baym et al., 2015) using the Nextera XT library kit (Illumina, San Diego, CA, USA). Quality and quantity controls of the extracted DNA and the DNA library were performed with Qubit (Thermo Fisher, USA) and Fragment Analyzer (Agilent Technologies, CA, USA). We used an Illumina NextSeq 2000 to sequence 2x150 bp paired-end reads, resulting in 40-60 M reads per sample, *i.e*., 6-9 Gb per sample. Subsequently, we performed human read removal with bbmap (minid=0.6 k=14 usemodulo bwr=0.16 fast minhits=1 qtrim=lr trimq=20 untrim) using the masked human genome HG19 (BBMap, 2016). Fastp [18] was applied for quality trimming and deduplication (--average_qual 30 –length_required 100 –trim_front1 10 –trim_tail1 4 –dedup), and forward and reverse reads were concatenated for subsequent analysis. The resulting datasets contained 0.65-1.25 M high-quality reads, *i.e.*, 98-188 Mb per sample, each with an estimated proportion of 20-36% microbial reads, as determined by Bracken v2.9 [19]. Metaphlan4 [20] as used to profile the microbial composition, and hclust2 (GitHub/Hclust2, 2020) was used to generate heatmaps based on filtered relative abundance tables (taxa represented with >0.1% relative abundance). To identify individual species in BALF specimens with DNA concentrations >1 ng/µL, we determined species abundances with Metalan4 and Bracken v2.9.

**Supplementary Results**

**Analysis of Bray-Curtis, and Jaccard indices at 98% OTUs level**

Analysis of Bray-Curtis, and Jaccard indices at the 98% OTUs indicated disease state remained significant in the presence-absence datasets: i) *adonis*: Bray-Curtis disease status R^2^ =0.0279, p =0.2386, age category R^2^ =0.2034, p =0.1693; Jaccard disease status R^2^ =0.05, p =0.0449, age category R^2^ =0.174, p =0.3356, based on 10^5^ permutations; ii) Constrained principal coordinates analysis: Constrained inertia, Bray-Curtis=0.2279, Jaccard=0.4999; “anova.cca” by term, disease status: explained variance, Bray-Curtis= 2.76%, p=0.2606, Jaccard=7.65%, p=0.057; age category: explained variance : Bray-Curtis=20.03%, p=0.1932, Jaccard=34.50%, p =0.40, based on 10^5^ permutations.

**Analysis of dispersion of microbiome structure within disease states**

To assess the inter-individual variability of the microbiome structure within each patient group, we performed the multivariate homogeneity of group dispersion with “betadisper” (vegan) using centroid parameter for Bray-Curtis, and Jaccard indices. Overall, dispersion within TB patients is higher compared to NTM patients, marginally significant in Bray-Curtis, coherent with our previous findings of indicator taxa whereby fractions of TB patients carry the indicator taxa which are hardly detected in NTM patients. Average distances to centroid were as follows: Bray-Curtis (TB =0.6368, NTM=0.5566), anova: p=0.09; Jaccard (TB=0.6037, NTM=0.5736), anova: p= 0.4989

**Analysis of identified indicator taxa**

To further inspect the identified indicator taxa of disease state, namely ASV_7, ASV_21, and OTU_11_0.02, for each indicator taxon, we constructed a linear model (lm) in R with relative abundances a response variable, and disease group, sex, age category, and sampling year as explanatory variables. When needed, square-root transformation was applied on relative abundances. Models were checked, and validated by i) Checking the distribution of the residuals, ii) plotting the fitted- against residual values, and iii) plotting the residuals against the response variables. The main linear model outputs are: ASV_7: summary lm: F_20, 21_=, R^2^_adj._= 0.427; p =0.02, ASV_21: summary lm: F_20, 21_=, R^2^_adj._= 0.1267; p =0.2791, OTU_11_0.02: summary lm: F_20, 21_=, R^2^_adj._=0.1267; p =0.1267. Proportions of variance explained by the disease status, age category, and sampling year are provided in Table 1.

**Table 1**. Analysis of variance of linear models fitted for ASV_7, ASV_21, and 0.02_OTU_11

| **Response variable** | **Explanatory variable** | **p value** | **Variance explained (%)** |
| --- | --- | --- | --- |
| **ASV_7** | Disease status | 0.0028 | 16 |
|  | Age category | 0.0306 | 27.65 |
|  | Sampling year | 0.1633 | 27 |
| **ASV_21** | Disease status | 0.0107 | 16.70 |
|  | Age category | 0.4948 | 14.07 |
|  | Sampling year | 0.5135 | 24.50 |
| **0.02_OTU_11** | Disease status | 0.0107 | 16.70 |
|  | Age category | 0.4948 | 14.07 |
|  | Sampling year | 0.51358 | 24.50 |

**Interactions between *Serratia* ASVs and OTUs within disease states**

Given the dominance and significance of unclassified *Yersiniaceae* and *Serratia* traits as disease state indicators, we aimed to explore the interactions between the abundances of the major *Yersiniaceae- and Serratia*-adjusted ASVs and 98% OTUs within TB, and NTM patients, distinctively. Among the TB patients, ASV_14 *Serratia* negatively correlated with ASV_1 unclassified *Yersiniaceae,* whereas ASV_14 Serratia positively correlated with ASV_3 unclassified *Yersiniaceae* (supplementary figure S4a and Supplementary Table 4). In NTM patients, ASV_14 *Serratia* also positively correlated with ASV_3 unclassified *Yersiniaceae,* while ASV_21 *Serratia* positively correlated with ASV_7 unclassified *Yersiniaceae* (supplementary figure 4b and supplementary table S4).

At the 98% OTUs level, and among TB patients, we observed a moderately strong negative correlation between the abundances of 0.02_OTU_1 unclassified *Yersiniaceae* and both *Serratia* OTUs (0.02_OTU_7 and 0.02_OTU_11). We found no significant correlations between major 98% OTUs within NTM patients (supplementary figure S5a, and S5b and supplementary table S4).

This striking result indicates when the ASVs are clustered to one higher resolution unit, the dominant traits likely drive the observed distinct interactions

**Verification of the clustering of adjusted ASVs, and 98% OTUs**

To further confirm the accuracy of the adjusted clustering of ASVs, and 98% OTUs, we aligned the following retrieved 16S rRNA gene sequences of *Serratia*, and its phylogenetically closest genus *Yersinia*: NR_041833.1 *Yersinia ruckeri* strain ATCC 29473, NR_115973.1 *Yersinia kristensenii* strain CCUG 11294, NR_042485.1 *Yersinia similis* strain Y228, NR_115976.1 *Yersinia ruckeri* strain CCUG 14190, NR_179466.1 *Yersinia alsatica* strain IP38850, NR_025340.1 *Serratia grimesii* strain DSM 30063, , NR_037112.1 *Serratia quinivorans* strain 4364, NR_042062.1 *Serratia liquefaciens* strain CIP 103238, NR_113616.1 *Serratia grimesii* strain NBRC 13537, NR_114575.1 *Serratia quinivorans* strain LMG 7887, NR_114576.1 *Serratia grimesii* strain LMG 7883, NR_121703.1 *Serratia liquefaciens* strain ATCC 27592, NR_121742.1 *Yersinia similis* strain Y228, NR_178702.1 *Serratia myotis* strain 12; along with the main unclassified *Yersiniaceae*/*Serratia* ASV sequences in Geneious Prime using Geneious alignment (global alignment with free end gaps & 93% similarity). After alignment was trimmed to equal length; based on the resulting similarity distance matrix, two distinct clusters of ASVs emerge: Cluster 1 includes six ASVs: ASV_1, ASV_3, ASV_7, ASV_11, ASV_12, ASV_22, and Cluster 2 includes four ASVs: ASV_4, ASV_14, ASV_21, ASV_25. The within ASVs-cluster similarity scores are between 99.358% and 99.786%, and the between cluster similarity scores are 97.877% and 98.093% (supplementary table S5). Interestingly, all ASVs of Cluster 1 classify as unclassified *Yersiniaceae*, while all ASVs of Cluster 2 classify as *Serratia*, and the BLAST algorithm (Camacho et al., 2009) suggest *S. grimeisi* / *S. liquefaciens* species for cluster 1 ASVs, and *S. quinivorans* / *S. myotis* for cluster 2 ASVs. Besides, we contrasted the two ASV clusters defined above with the 98% OTUs defined based on the alignment in DECIPHER (2.24.0) of the entire dataset. We find that these two methods disagree on the clustering of two ASVs: ASV_4 which was clustered to OTU_1_0.02, but to Cluster 2 based on the Geneious. Also, ASV_21 clustered apart to OTU_11_0.02 while based Geneious it belongs to cluster 2 (see below):

**98% clustering based on alignment in DECIPHER**

**OTU_1_0.02**: 07/10 ASVs, ASV_1, ASV_4, ASV_7, ASV_3, ASV_11, ASV_12, ASV_22

**OTU_7_0.02**: 02/10 ASVs, ASV_14, ASV_ 25

**OTU_11_0.02**: 01/10 ASVs, ASV_21

**Geneious based clustering**

**Cluster 1**: 06/10 ASVs, ASV_1, ASV_3, ASV_7, ASV_11, ASV_12, ASV_22

**Cluster 2**: 04/10 ASVs, ASV_4, ASV_14, ASV_21, ASV_25

The distance between the two clusters ranges from 97.452% to 97.877% similarity.

**Exploratory assessment of *Serratia* taxonomic profiles**

The relative abundances of the *Yersiniciaceae/Serratia* taxa were comparable to the 16S rRNA data for three selected specimens from patients with NTM lung disease. At the species level, metaphlan4 indicated the presence of *Serratia grimesii* (supplementary figure S6), while Bracken consistently reported multiple *Serratia* species with similar relative abundances. However, the same *Serratia*-specific abundance profile was observed when analyzing a *Serratia grimesii* type strain with Bracken, suggesting limitations with the associated taxonomic database for the genus *Serratia.* Additionally, using the BLAST algorithm [21] on the amplicon short reads representing the above highlighted ASVs of *Serratia* and “unclassified *Yersiniaceae*” supported the metaphlan4 results for the selected specimens. ASV_1, ASV_3, and ASV_7 had the highest sequence similarity scores with those of *Serratia liquefaciens* and *Serratia grimesii*. ASV_4, ASV_14, and ASV_21 had the highest sequence similarity scores for *Serratia myotis* and *Serratia quinivorans*. These exploratory data indicate that some key traits in our analysis were distinct for *Serratia grimesii sub-species/*strains; however, we cannot exclude the presence of other *Serratia* species, especially low-abundance species and strains.
